# Gut microbiota regulates food intake in a rodent model of binge-eating disorder

**DOI:** 10.1101/2024.07.05.602243

**Authors:** Thomas Demangeat, Léa Loison, Marion Huré, do Rego Jean-Luc, Pierre Déchelotte, Najate Achamrah, Moïse Coëffier, David Ribet

## Abstract

**Objective:** Binge-eating disorder is characterized by recurrent episodes of consumption of large amounts of food within a short period of time, without compensatory behaviours. This disease is a major public health issue since it decreases patients quality of life and is associated with numerous comorbidities, encompassing anxiety, depression and complications associated with obesity. The pathophysiology of binge-eating disorder is complex and involves both endogenous, environmental and sociocultural factors. The gut microbiota has been proposed to be an important player in the onset or maintenance of eating disorders. Here, we aim to better delineate the potential role of the gut microbiota in binge-eating disorder.

**Method:** We used a model of binge-eating disorder where eight-weeks-old C57Bl/6 female mice had access during 2 hours, every 2 days over a 10-day period, to a highly palatable and high-calorie diet. Half of the animals received antibiotics to deplete their gut microbiota. Eating behaviour and other behavioural parameters were compared between groups.

**Results:** We observed an increase in food intake in mouse exposed to high-fat high-sucrose diet, as well as tachyphagia and craving for food during binge-eating episodes. We demonstrate the gut microbiota depletion further increases food intake, specifically during binge-eating episodes.

**Discussion:** These results show that the gut microbiota is involved in the control of food intake during episodes of binge-eating. This strengthens the potential role of the gut bacteria in binge-eating disorder and open the way for future therapeutic strategies aiming at targeting patients’ gut microbiota.

## Introduction

Binge-Eating Disorder (BED) is an eating disorder marked by recurrent episodes of consumption of large amounts of food within a short period of time (often less than two hours) (1). These episodes are associated with an overwhelming sense of loss of control over eating. In contrast to bulimia nervosa, inviduals with BED do not perform inappropriate compensatory behaviors (*e.g*., purging, fasting or excessive exercising). In accordance with DSM-5 criteria, BED’s associated binge-eating episodes should occur at least one day a week for 3 months and are frequently associated with fast eating, eating until feeling uncomfortably full, eating without feeling physically hungry, eating alone, or feeling disgusted with oneself or guilty after overeating (2). The incidence of BED has strongly increased in recent years, and more particularly since the COVID-19 pandemic (3). The estimated lifetime prevalence of BED is approximately 2.8% in women and 1% in men (4).

BED is closely linked to numerous comorbidities, encompassing anxiety, depression, functional gastrointestinal disorders, and complications associated with obesity, including metabolic disorders (5). This disease exhibits striking parallels with addictive behavior and is frequently subjected to social stigma (6, 7). As such, BED is strongly associated with distress and a reduced quality of life.

The current treatment of BED consists in regular monitoring by dietitians and nutritionists, psychotherapy, adapted physical activity, and, in some cases, the use of serotonin reuptake inhibitors (8–10). Despite these different approaches, the success of current therapies remains limited and the relapse rate still exceed 30% at the ten-year mark (11). This is partly due to the poor understanding of the pathophysiological mechanisms underlying BED. Factors contributing to the pathophysiology of BED encompass psychological, sociocultural, and genetic factors (12,13). Additional factors, such as the gut microbiota, have recently been proposed as important players in the etiology of the disease (14). Indeed, the gut microbiota plays critical roles in the regulation of eating behavior, in energy metabolism as well as in the development of anxiety, depression and functional digestive disorders (15,16). Thus, the gut microbiota may contribute to the onset and/or maintenance of BED. A gut dysbiosis has been described in BED patients, although the number of clinical studies focusing on the gut microbiota composition in this disease remains very limited (17,18). Yet, the functional consequences of this gut dysbiosis in BED pathophysiology remains unknown and further work is required to establish causal relationships.

The use of experimental animal models is pivotal to unveil the pathophysiological mechanisms of BED. Several animal models of BED are described in the scientific literature (19). Most of these models are based on the exposure of rodents to high-calorie and/or highly palatable diets for a limited duration (generally less than 2 hours), in order to trigger episodes of intense food consumption mirroring those observed in BED patients. We aimed to use one of this BED murine model to explore the potential contribution of the gut microbiota in eating behaviour and comorbidities associated with this eating disorder.

## Methods

### Animals

Animal care and experimentation were approved by a regional Animal Experimentation Ethics Committee (APAFIS #38597-2022091914519369 v3) and complied with the guidelines of the European Commission for the handling of laboratory animals (Directive 2010/63/EU). All efforts were made to minimize suffering of animals.

Eight-weeks-old C57Bl/6 female mice (Janvier Labs, Le-Genest-Saint-Isle, France) were first housed at 23°C in regular open cages with a 12h light-dark cycle for seven days in a ventilated cabinet. Independent cages of male mice were introduced in the same cabinet to promote the synchronization of the estrous cycles in females. Half of the female animals were treated during this first week with both antifungals and antibiotics, diluted in drinking water, to trigger gut microbiota depletion (20). For this, animals had first access to drinking water with Amphotericin-B (Sigma-Aldrich) for 3 days (0.005 mg/mL), and then to drinking water containing Amphotericin-B (0.005 mg/mL), Ampicillin (0.5 mg/mL, Sigma-Aldrich), Neomycin trisulfate salt hydrate (0.5 mg/mL, Sigma-Aldrich), Metronidazole (0.5 mg/mL, Sigma-Aldrich) and Vancomycin hydrochloride (0.25 mg/mL, Sigma-Aldrich) (20). Drinking water was renewed every day. After this first week, female mice were individually housed at 23°C with a 12h light-dark cycle in BioDAQ cages (Research Diets Inc.), to allow for a continuous monitoring of eating behaviour. Administration of antifungals and antibiotics was continued for half of the animals. Each animal was assigned to one of the following group (8 animals/group): a control group (CTRL group), a group periodically exposed to a high fat/high sucrose (HFHS) diet (HFHS group), a group treated with antibiotics (CTRL+ATB group) and a group both periodically exposed to a HFHS diet and treated with antibiotics (HFHS+ATB group) (Fig. S1). Allocation to the different groups was performed in order to minimize body weight variations between groups.

Mice from the HFHS and HFHS+ATB groups had access during 2 hours (at the beginning of the dark phase), every 2 days over a 10-day period, to a diet enriched in carbohydrates and lipids (HFHS diet, D12451i, Research Diets Inc.) (Fig. 1). This diet contains 45 kcal % fat and 21 mass % sucrose. These short and repeated exposures to an HFHS diet aims to mimic episodes of binge eating. Between these episodes, animals from the HFHS and HFHS+ATB groups had an *ad libitum* access to a standard diet (Teklad Global 16% Protein Rodent Diet, INOTIV®). Mice from the CTRL and CTRL+ATB groups had an *ad libitum* access to a standard diet during the whole protocol.

**Fig 1:**
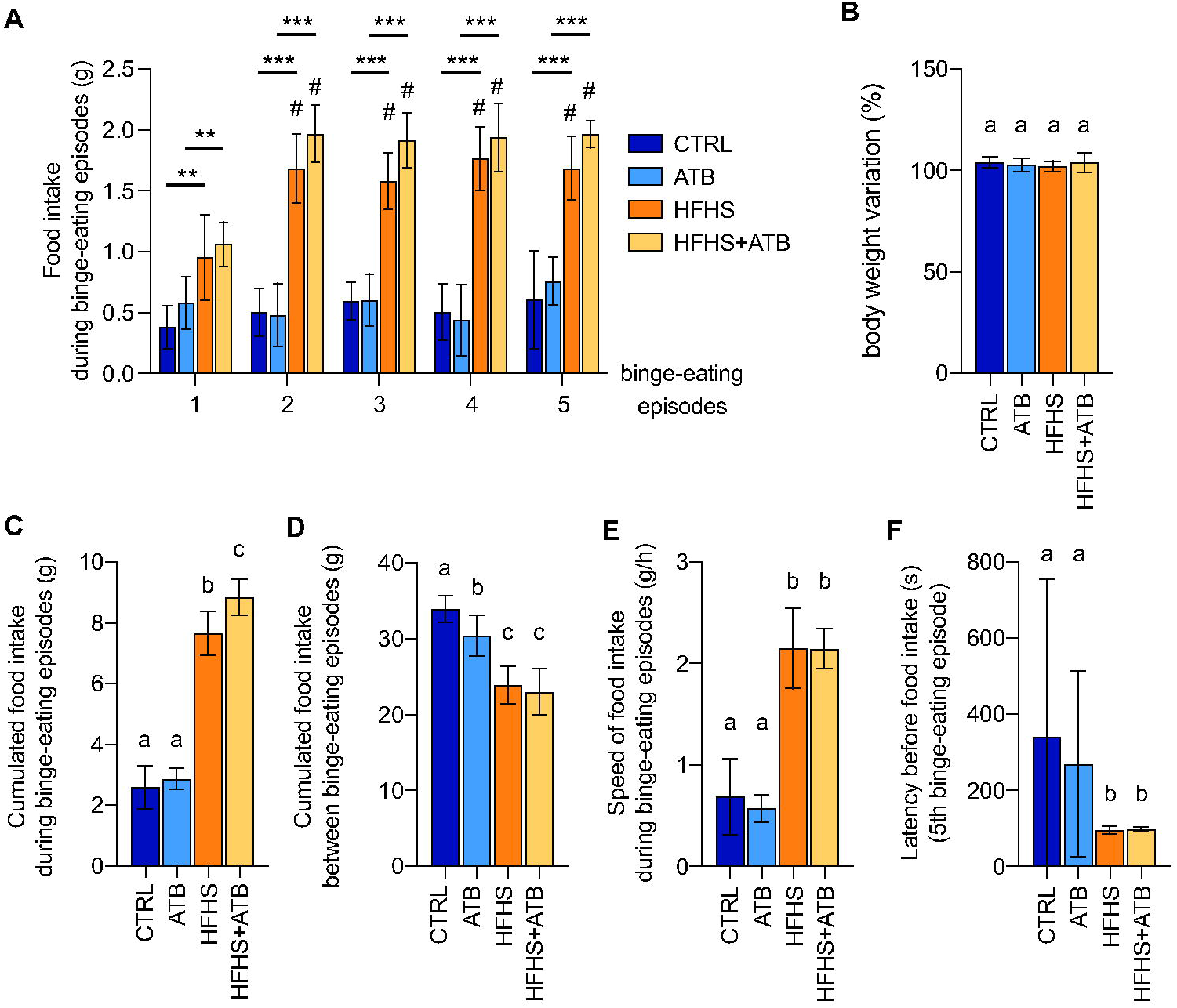
Gut microbiota-dependent modulation of eating behaviour in a rodent BED model. (A) Food intake in each of the five binge-eating episodes (mean ± s.d.; n= 8/group; 2-way ANOVA; **, p<0.01; ***, p<0.001; #, p<0.05 vs the first episode). (B) Body weight variation at the end of the protocol (percentage of initial body weight) (mean± s.d.; n= 8/group; 1-way ANOVA with Tukey’s correction). (C-D) Cumulated food intake during (C) or between (D) binge-eating episodes (mean± s.d.; n= 8/group; 1-way ANOVA with Tukey’s correction). **(E)** Speed of food intake during binge-eating episodes (mean ± s.d.; n= 8/group; 1-way ANOVA with Tukey’s correction). (F) Latency between the access to **HFHS** diet and the first event of food intake (mean ± s.d.; n= 8/group; Kruskal-Wallis test with Dunn’s correction). Labeled means without a common letter differ. Similar results were observed for 2 independant animal series.

Body weight was monitored daily, and body composition was assessed on vigil animals at the end of the protocol using fast nuclear magnetic resonance (Minispec LF110, Brucker). To evaluate depressive-like behaviour, mice were subjected to a nesting test, 2 days after the last binge-eating episode. In this test, a compressed cotton nestlet is placed in cages and a score is given to the nest constructed by mice from this cotton nestlet after 24 hours (21) (Fig. S1). To evaluate anxiety-like behaviour, mice were subjected to a light-dark compartment test 3 days after the last binge-eating episode (21) (Fig. S1).

At the end of the protocol, all animals were euthanized. Cecal contents were collected and stored at -80°C until analysis. Two independent animal series were performed.

### Quantification of cecal bacterial densities

DNAs from mouse cecal contents were extracted using the QIAamp DNA Stool Mini Kit (QIAGEN), including a bead-beating step (0.1 mm zirconia silica beads, BioSpec products), as previsouly described (22). Quantitative real-time polymerase chain reaction (qPCR) was performed on DNA samples contents to monitor the efficiency of antibiotics-mediated gut bacterial depletion. To quantify total Eubacteria, qPCR were performed using Itaq Universal SYBR Green Supermix and primers targeting conserved regions in the 16S rRNA gene (Eub-338 F, 5’-ACTCCTACGGGAGGCAGCAG-3’ and Eub-518 R, 5’-ATTACCGCGGCTGCTGG-3’) (23). The Cq determined in each sample were compared with a standard curve made by diluting genomic DNA extracted from a pure culture of *E. coli*, for which cell counts were determined prior to DNA isolation.

## Results

We used a previously described BED rodent model in which animals were exposed every 2 days over a 10-day period to a diet enriched in carbohydrates and lipids (high fat-high sucrose (HFHS) diet; Fig. S1A). These short and repeated exposures to an HFHS diet aims to mimic episodes of binge eating. Between these episodes, animals had an *ad libitum* access to a standard diet (19). In order to delineate the potential role of the gut microbiota in eating behaviour in this BED model, we treated mice with antibiotics in order to deplete their gut microbiota (20). The efficiency of depletion was validated by quantifying the density of Eubacteria in mouse cecal contents (Fig. S1B). Four groups of mice were compared in this study: a control group (CTRL group), a group periodically exposed to HFHS diet (HFHS group), a group treated with antibiotics (CTRL+ATB group) and a group both periodically exposed to HFHS diet and treated with antibiotics (HFHS+ATB group) (Fig. S1A).

We first monitored food intake in the different groups. We observed a significant increase in food intake for the HFHS and HFHS+ATB groups, compared to the CTRL and ATB groups, respectively, for each episode of access to the HFHS diet (Fig. 1). Of note, the food intake during these episodes increases in HFHS and HFHS+ATB groups between the first episode and the following episodes. We also observed that the average speed of food intake is significantly increased in animals exposed to the HFHS diet compared to control animals (Fig. 1). These increases in food intake and in speed of eating mimic the binge-eating episodes and the tachyphagia classically observed in BED patients. The latency between the opening of the HFHS food access and the first food intake event is significantly decreased compared to animals fed with regular diet. This probably reflects an increase in craving for highly palatable food triggered along the protocol. Between “binge-eating episodes”, animals from the HFHS and HFHS+ATB groups exhibit reduced food intake (Fig. 1). These results suggest the presence of compensatory behaviour following binge-eating episodes. Accordingly, we did not observe any significant difference in body weight or body composition between CTRL and HFHS groups at the end of the protocol, despite the consumption of a diet rich in calories.

To assess the impact of the gut microbiota in our BED rodent model, we compared food intakes between the HFHS and HFHS+ATB groups. Interestingly, we observed a significant increase in the cumulated food intake of the HFHS+ATB group compared to the HFHS group during the binge-eating episodes, but not between episodes (Fig. 1C and 1D). These results suggest that the depletion of the gut microbiota modifies mouse eating behaviour during binge-eating episodes. This depletion does not affect the speed of food intake nor the craving of animals for the HFHS diet.

Similar results were observed for two independent animal series.

We then evaluated if the BED model used in this study triggers anxiety-or depressive-like disorders in animals. To evaluate depressive-like behaviours, we performed a nesting test, 2 days after the last binge-eating episode, by providing a compressed cotton nestlet to the animals and by scoring the structure of the nest built by animals after 24 hours (21). A poor nest score indicates depressive-like behaviours. No significant differences in nest scores were observed between groups (Fig. 2A). To evaluate anxiety, animals were placed in light/dark boxes 3 days after the last binge-eating episode and the time spent in each compartment during the 15 first minutes of the test was recorded. The light compartment constitutes an aversive environment for rodents in this test. As such, a decrease in the time spent in this compartment is defined as an anxiety-like behaviour (21). The time spent in the light compartment was not significantly different between groups (Fig. 2B). Together, these results suggest that the BED model used in this study does not trigger anxiety-or depressive-like behaviours. Gut depletion has no further impact on animal behaviour.

**Fig 2:**
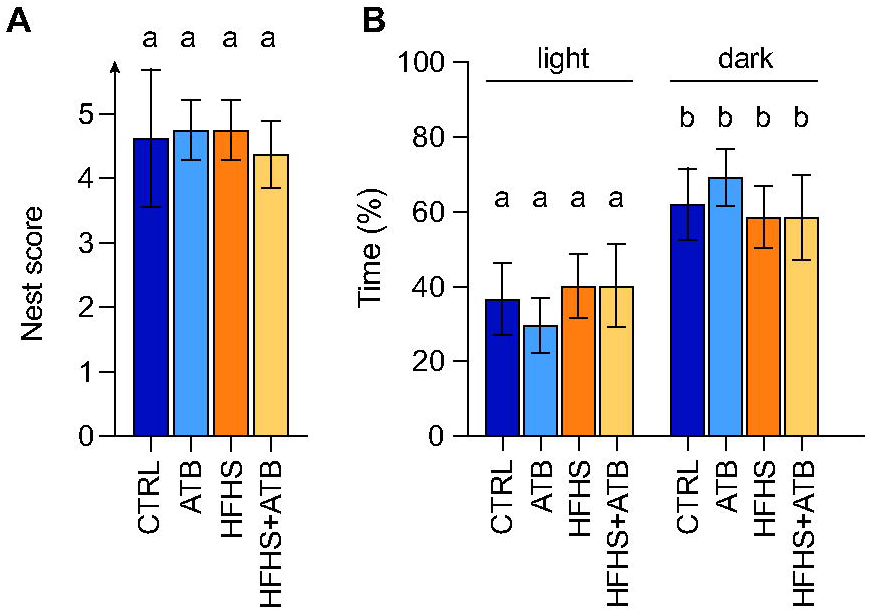
Lack of anxiety- and depressive-like disorders triggered in animals exposed to a BED model. (A) Nest score from the nesting test (mean ± s.d.; n= 8/group; 1-way ANOVA with Tukey’s correction). Labeled means without a common letter differ. Similar results were observed for 2 independant animal series. (B) Time spent in light or dark compartments during the light/dark box test (mean ± s.d.; n= 8/group; 1-way ANOVA with Tukey’s correction). Similar results were observed for 2 independant animal series.

## Discussion

Medical care for patients with Binge-Eating Disorder is a real challenge today. Animal models are valuable tools for studying the pathophysiology of this disease and for deciphering the role of complex inter-organ interactions, such as the microbiota-gut-brain axis. In order to better delineate the potential role of the gut microbiota in BED, we used here a model based on intermittent accesses to a highly palatable and high-calorie containing food. We used mice since they usually better respond to BED models than rats, and focused on females, to mirror the female predominance of this disease (19).

Our model recapitulates several important features of BED models such as an escalation of food intake between the beginning and the end of the protocol in the BED groups, tachyphagia and an observed craving for food during binge-eating episodes. Interestingly, between episodes, mice decreased food consumption, suggesting that compensatory behavior are occuring, which align with patients behaviour exhibiting moderate restriction to offset excessive caloric consumption, often driven by post-meal guilt, ultimately culminating in a loss of control over eating (24). This BED model is distinct from an obesity model since no increase in body weight was observed. Our model does not recapitulate some of the classical comorbidities associated with BED such as anxiety or depression. This might be due to the short duration of our model, the lack of alternance with periods of food restriction or the lack of stress (such as foot-shock stress) in our experimental design, which are sometimes used in other BED models (19).

We demonstrate that depletion of the gut microbiota promotes food intake in our BED model. Thus, the gut microbiota plays a role in food intake control, particularly in curtailing calorie consumption during binge-eating episodes. This result strongly strengthens the hypothesis that alteration of the gut microbiota, as observed in BED patients, may favor loss of control in food consumption and promote the altered eating behaviour observed in BED.

Interestingly, it has been reported that antibiotic depletion of the gut microbiota in mice results in overconsumption of highly palatable food such as high-sucrose pellets (25). Another study suggested that the gut microbiota is involved in overeating disorders, where patients present with experience of cravings for palatable food (26). In this study, a mouse model combining stress with a history of dieting was associated with a dysbiosis and an alteration of the gut metabolome, which ultimately results in an alteration of the gut-brain axis and an hedonic overconsumption of highly palatable food (26).

Altogether, these results demonstrate the central role of the gut microbiota in consumption of highly palatable food and binge-eating behaviour. They pave the way for the better understanding of the molecular determinants of the cross-talk between gut bacteria and their host and for the design of future therapeutic strategies aiming at manipulating patients’ gut microbiota to improve the management of eating disorders.

## Public Significance

Binge-Eating Disorder (BED) is a widespread yet challenging condition to treat. This disease is characterized by recurrent episodes of consumption of large amounts of food within a short period of time. The mechanisms underlying this disease are not fully understood. It has recently been proposed that the gut microbiota plays a significant role in the onset of this disease since gut bacteria can communicate with the central nervous system and regulate food intake. In this study, we evaluated the role of the gut microbiota in BED using a dedicated mouse model. Our results show that the gut microbiota regulates food intake during binge-eating episodes. They strengthen the hypothesis that an alteration of the gut microbiota composition may be a potential susceptibility factor for BED. This work opens new perspectives for the development of innovative treatments based on microbiota modulation that could improve the management of this eating disorder.

## Data availability statement

The authors confirm that the data supporting the findings of this study are available within the article and its supplementary materials.

## Figures

**Figure S1: BED model and antibiotic-mediated gut microbiota depletion**.

(A) Experimental protocol. (B) Quantification of Eubacteria in mouse cecal contents (mean ± s.d.; n=8/group; Kruskal-Wallis test with Dunn’s correction). Labeled means without a common letter differ. Similar results were observed for 2 independant animal series.

## REFERENCES

1. Wonderlich SA, Gordon KH, Mitchell JE, Crosby RD, Engel SG. The validity and clinical utility of binge eating disorder. Int J Eat Disord. 2009;42(8):687□705.

2. Diagnostic and statistical manual of mental disorders: DSM-5™, 5th ed. Arlington, VA, US: American Psychiatric Publishing, Inc.; 2013. xliv, 947 p. (Diagnostic and statistical manual of mental disorders: DSM-5™, 5th ed).

3. Tavolacci MP, Ladner J, Dechelotte P. COVID-19 Pandemic and Eating Disorders among University Students. Nutrients. 2021;13(12):4294.

4. Galmiche M, Déchelotte P, Lambert G, Tavolacci MP. Prevalence of eating disorders over the 2000-2018 period: a systematic literature review. Am J Clin Nutr. 2019;109(5):1402□13.

5. Mitchell JE. Medical comorbidity and medical complications associated with binge-eating disorder. Int J Eat Disord. 2016;49(3):319□23.

6. Gearhardt AN, White MA, Masheb RM, Morgan PT, Crosby RD, Grilo CM. An examination of the food addiction construct in obese patients with binge eating disorder. Int J Eat Disord. 2012;45(5):657□63.

7. Ebneter DS, Latner JD. Stigmatizing attitudes differ across mental health disorders: a comparison of stigma across eating disorders, obesity, and major depressive disorder. J Nerv Ment Dis. 2013;201(4):281□5.

8. Linardon J, Wade TD, de la Piedad Garcia X, Brennan L. The efficacy of cognitive-behavioral therapy for eating disorders: A systematic review and meta-analysis. J Consult Clin Psychol. 2017;85(11):1080□94.

9. Raisi A, Zerbini V, Piva T, Belvederi Murri M, Menegatti E, Caruso L, et al. Treating Binge Eating Disorder With Physical Exercise: A Systematic Review and Meta-analysis. J Nutr Educ Behav. 2023;55(7):523□30.

10. Arnold LM, McElroy SL, Hudson JI, Welge JA, Bennett AJ, Keck PE. A placebo-controlled, randomized trial of fluoxetine in the treatment of binge-eating disorder. J Clin Psychiatry. 2002;63(11):1028□33.

11. Safer DL, Lively TJ, Telch CF, Agras WS. Predictors of relapse following successful dialectical behavior therapy for binge eating disorder. Int J Eat Disord. 2002;32(2):155□63.

12. Javaras KN, Laird NM, Reichborn-Kjennerud T, Bulik CM, Pope Jr HG, Hudson JI. Familiality and heritability of binge eating disorder: Results of a case-control family study and a twin study. Int J Eat Disord. 2008;41(2):174□9.

13. Treasure J, Duarte TA, Schmidt U. Eating disorders. Lancet Lond Engl. 2020;395(10227):899□911.

14. Guo W, Xiong W. From gut microbiota to brain: implications on binge eating disorders. Gut Microbes. 2024;16(1):2357177.

15. Margolis KG, Cryan JF, Mayer EA. The Microbiota-Gut-Brain Axis: From Motility to Mood. Gastroenterology. 2021;160(5):1486□501.

16. Han H, Yi B, Zhong R, Wang M, Zhang S, Ma J, et al. From gut microbiota to host appetite: gut microbiota-derived metabolites as key regulators. Microbiome. 2021;9(1):162.

17. Leyrolle Q, Cserjesi R, Mulders MDGH, Zamariola G, Hiel S, Gianfrancesco MA, et al. Specific gut microbial, biological, and psychiatric profiling related to binge eating disorders: A cross-sectional study in obese patients. Clin Nutr. 2021;40(4):2035□44.

18. Castellini G, Cassioli E, Vitali F, Rossi E, Dani C, Melani G, et al. Gut microbiota metabolites mediate the interplay between childhood maltreatment and psychopathology in patients with eating disorders. Sci Rep. 2023;13(1):11753.

19. Rehn S, Raymond JS, Boakes RA, Leenaars CHC. A systematic review and meta-analysis of animal models of binge eating - Part 1: Definitions and food/drink intake outcomes. Neurosci Biobehav Rev. 2022;132:1137□56.

20. Tirelle P, Breton J, Riou G, Déchelotte P, Coëffier M, Ribet D. Comparison of different modes of antibiotic delivery on gut microbiota depletion efficiency and body composition in mouse. BMC Microbiol. 2020;20(1):340.

21. Kremer M, Becker LJ, Barrot M, Yalcin I. How to study anxiety and depression in rodent models of chronic pain? Eur J Neurosci. 2021;53(1):236□70.

22. Breton J, Tirelle P, Hasanat S, Pernot A, L’Huillier C, do Rego JC, et al. Gut microbiota alteration in a mouse model of Anorexia Nervosa. Clin Nutr Edinb Scotl. 2021;40(1):181□9.

23. Fierer N, Jackson JA, Vilgalys R, Jackson RB. Assessment of Soil Microbial Community Structure by Use of Taxon-Specific Quantitative PCR Assays. Appl Environ Microbiol. 2005;71(7):4117□20.

24. Masheb RM, Grilo CM. On the Relation of Attempting to Lose Weight, Restraint, and Binge Eating in Outpatients with Binge Eating Disorder. Obes Res. 2000;8(9):638□45.

25. Ousey J, Boktor JC, Mazmanian SK. Gut microbiota suppress feeding induced by palatable foods. Curr Biol CB. 2023;33(1):147-157.e7.

26. Fan S, Guo W, Xiao D, Guan M, Liao T, Peng S, et al. Microbiota-gut-brain axis drives overeating disorders. Cell Metab. 2023;S1550-4131(23)00335-2.

