## Supplemental Figure 1 for "Gut microbiota regulates food intake in a rodent model of binge-eating disorder"

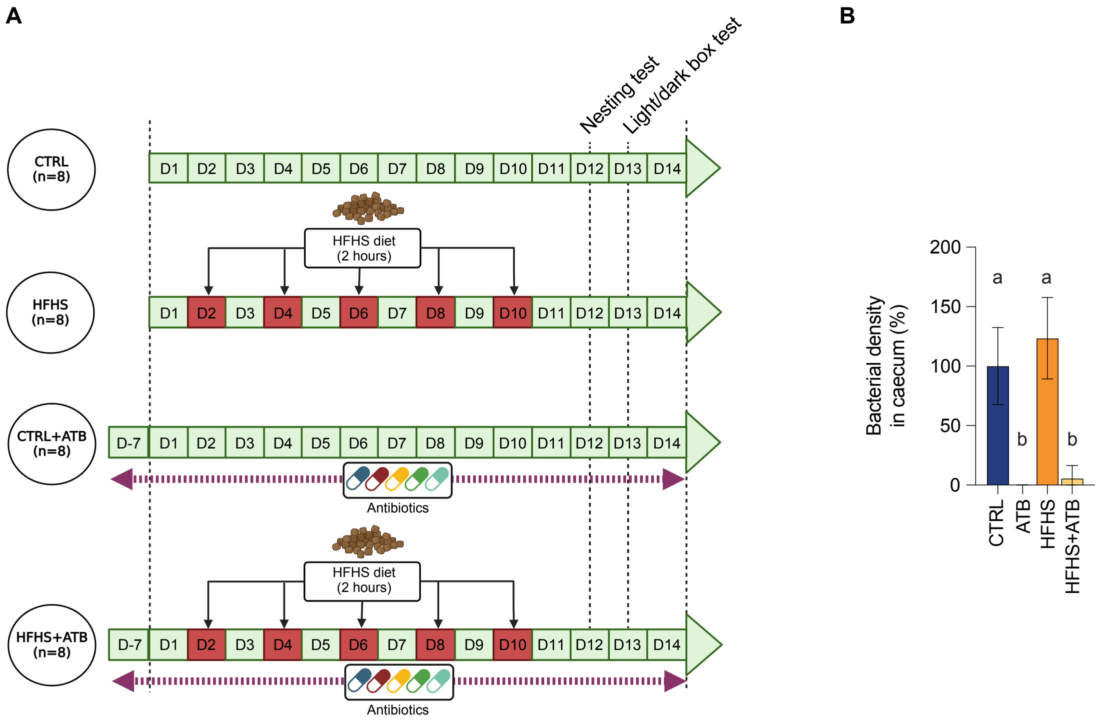


**Fig. S1: BED model and antibiotic-mediated gut microbiota depletion**

(A) Experimental protocol. (B) Quantification of Eubacteria in mouse cecal contents (mean ± s.d.; n=8/group; Kruskal-Wallis test with Dunn's correction). Labeled means without a common letter differ. Similar results were observed for 2 independant animal series.
